# Autophagy is responsible for the accumulation of proteogenic dipeptides in response to heat stress in *Arabidopsis thaliana*

**DOI:** 10.1101/2020.03.25.004978

**Authors:** Venkatesh P. Thirumalaikumar, Mateusz Wagner, Salma Balazadeh, Aleksandra Skirycz

## Abstract

Proteogenic dipeptides are intermediates of proteolysis as well as an emerging class of small-molecule regulators with diverse and often dipeptide-specific functions. Herein, prompted by differential accumulation of dipeptides in a high-density *Arabidopsis thaliana* time-course stress experiment, we decided to pursue an identity of the proteolytic pathway responsible for the buildup of dipeptides under heat conditions. By querying dipeptide accumulation versus available transcript data, autophagy emerged as a top hit. To examine whether autophagy indeed contributes to the accumulation of dipeptides measured in response to heat stress, we characterized the loss-of-function mutants of crucial autophagy proteins to test whether interfering with autophagy would affect dipeptide accumulation in response to the heat treatment. This was indeed the case. This work implicates the involvement of autophagy in the accumulation of proteogenic dipeptides in response to heat stress in Arabidopsis.

## Introduction

Chemically speaking, dipeptides are a pair of amino acids joined by a single peptide bond. Considering only the combination of the 20 proteogenic amino acids, discussed here, the total number of dipeptides is 400. In the past, dipeptides have been exclusively considered as short-lived intermediates on their way to specific amino acid degradation pathways following further proteolysis. However, this view has been challenged by recent studies reporting regulatory roles for various proteogenic dipeptides across different organisms.

A compelling example comes from the regulation of the yeast (*Saccharomyces cerevisiae*) N-end rule pathway. UBR1 (Ubiquitin Protein Ligase E3 Component N-Recognin 1) is an ubiquitin ligase that has three ligand binding sites, of which two are specific for dipeptides. The consecutive binding of a dipeptide with the basic N-terminal residue (Arg, Lys, or His) to site-1 of UBR1 and of a dipeptide with the bulky hydrophobic N-terminal residue to site-2 (Trp, Phe, Tyr, Leu, or Ile) enables binding and ubiquitylation of the CUP9 protein, a transcriptional repressor of peptide import [1]. Other regulatory dipeptides include Tyr-Arg, also known as kyotorphin, a neurotransmitter, and a potent analgesic [2]. Interestingly, kyotorphin is produced in cells via active synthesis mediated by the enzyme kyotorphin synthase [3]. A different Tyr-containing dipeptide, Tyr-Leu, was demonstrated to display anxiolytic (i.e., stress-reducing) activity in mice, its potency being comparable to that of valium [4]. Notably, both the retro-sequence peptide Leu-Tyr and the mixture of tyrosine and leucine were inactive, demonstrating that even chemically similar dipeptides can have specific bioactivities. Moreover, the lack of anxiolytic activity reported for the amino acids supported the notion that the dipeptide function would be independent of the degradation to the constituent amino acids. In other words, when it comes to function, dipeptides are more than a sum of the constituent amino acids [4].

Recently, to identify potential regulatory small molecules on account of their presence in protein complexes, we developed a simple approach based on co-fractionation that exploits a significant size difference between protein□metabolite complexes and free metabolites [5] [6]. Remarkably, among the small molecules co-fractionating with proteins and protein complexes isolated from a native *Arabidopsis thaliana* lysate, dipeptides represented a large and diverse group. Specifically, of the 237 detected proteogenic dipeptides, 106 were classified as protein-bound; dipeptide elution profiles span the whole protein separation range, indicating the presence of multiple protein-binding partners. By focusing on one selected dipeptide, namely, Tyr-Asp, we could narrow down the list of Tyr-Asp interactors to the glycolytic enzyme glyceraldehyde-3-phosphate dehydrogenase (GAPDH). By testing multiple dipeptides and multiple GAPDHs, we demonstrated that the binding is specific to Tyr-Asp for both plant and mammalian GAPDHs [6]. In addition to its enzymatic function, GAPDH plays additional roles in non-metabolic processes such as the control of gene expression and actin-bundling [7]. However, the exact biological function of the Tyr-Asp–GAPDH interaction remains unclear; specific binding of the Tyr-Asp to GAPDH provides supporting evidence to the previously reported regulatory roles of the proteogenic dipeptides.

Intrigued by their presence in the Arabidopsis protein complexes, we decided to pursue the functions of the diverse proteogenic dipeptides. One immediate and essential question concerns dipeptides’ biogenesis. Proteolysis is considered the main source of the proteogenic dipeptides. Plant genomes encode multiple proteases with assigned exopeptidase activity. Exopeptidases would cleave amino acids but also dipeptides from either the C- (carboxypeptidases) or N-(aminopeptidases) termini of the protein substrates. Notably, known dipeptidyl exopeptidases are characterized by cleavage specificity, which could result in specific accumulation patterns. For instance, animal dipeptidyl peptidase-4 serine exopeptidase would cleave X-proline or X-alanine dipeptides from the N-terminus of peptides exclusively [8]. Dipeptides are subsequently cleaved to single amino acids by enzymes with dipeptidase activity. Similar to exopeptidases, dipeptidases also display substrate specificity [9].

Herein, prompted by the differential accumulation of proteogenic dipeptides in a high-density Arabidopsis time-course stress experiment [10,11], which we re-annotated using a dipeptide library, we decided to pursue an identity of the proteolytic pathway responsible for the increase of dipeptides measured under heat conditions. Co-expression analysis identified autophagy as a top hit, on par with previous reports demonstrating the importance of autophagy for plant response to stress [12]. During autophagy, cytoplasmic components including proteins are captured into a double-membraned structure, namely, the autophagosome, and delivered to the vacuole for recycling. The plant vacuolar proteome comprises multiple proteases, which subsequently cleave protein cargos to amino acids, which are thereafter transported back to the cytosol by amino acid transporters [13]. Notably, in the context of our findings, the vacuolar proteome is also characterized by the presence of dipeptidyl exopeptidases and dipeptide transporters [13]. In Arabidopsis, the role of autophagy has been studied extensively, benefiting from the multiple loss-of-function mutants as well as small-molecule inhibitors, which interfere with autophagosome formation, protein cargo recruitment, or vacuolar function [14]. Following accepted strategies, we used four autophagy mutants, to test whether disrupting autophagy would affect dipeptide accumulation in response to the heat treatment. Indeed, this was the case. In summary, we demonstrate that autophagy is involved in the accumulation of proteogenic dipeptides in response to heat stress.

## Results

### The levels of dipeptides’ response to varying light and temperature conditions

To gain insight into dipeptide biogenesis and function, we investigated dipeptide accumulation in the previously published Arabidopsis stress time-course experiment [10]. The experiment comprised eight different environmental conditions varying in temperature and light, and 22 time-points. All samples were subjected to transcriptomics [10] and untargeted metabolomics analysis [11]. Herein, we retrieved an existing metabolomics dataset [11] to annotate dipeptides by matching metabolic features, previously assigned as unknown, with a dipeptide reference compound library.

In total, we annotated 54 dipeptides, which were clustered into six distinctive groups **(Figure 1; Supplemental Table S1)**. Dipeptides in clusters 1 to 3 accumulated under high light conditions. Cluster 4 was mainly composed (16 of 21) of aspartic acid or glutamic acid-containing dipeptides (referred to later as acidic dipeptides) that responded rapidly to multiple stress conditions, with the levels increasing under heat and dark conditions but decreasing in the cold treatment. Dipeptides in clusters 5 and 6 were characterized by the presence of branch-chain amino acids (valine, leucine, isoleucine) and accumulated in all tested conditions except for high light, but the increase was moderate and measured 6 h or longer after the stress onset. Based on the clearly defined accumulation pattern, we decided to focus on the dipeptides of cluster 4.

**Figure 1.**
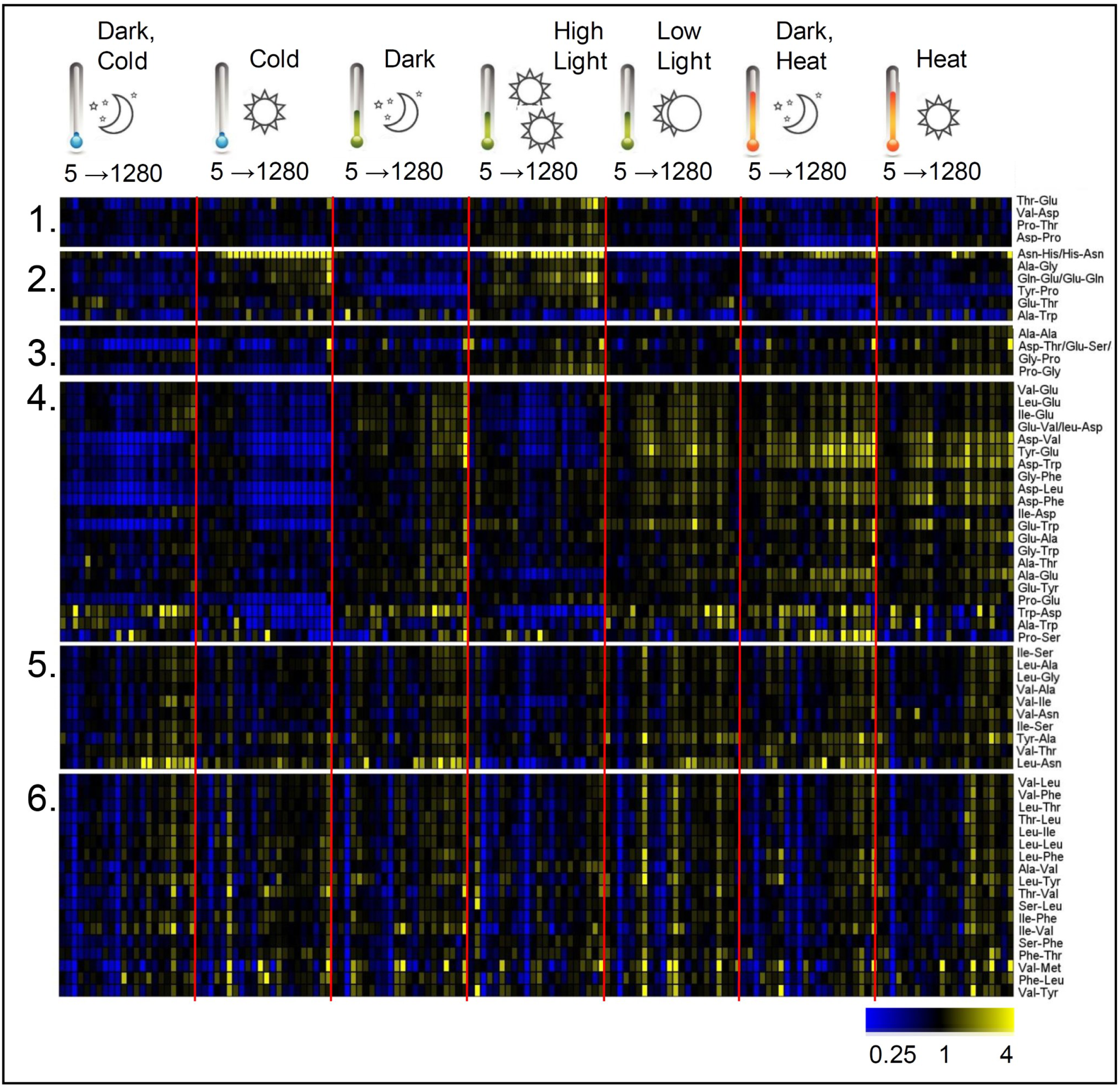
Heat map of dipeptide accumulation in the stress time series experiment. Data are expressed as fold change calculated using paired control treatment (21 °C and 150 μmol m^−2^s^−1^). Blue stands for down and yellow for upregulation. The figures 5 to 1280 refer to the first and last sampling time points given in minutes. Data were clustered using the Self Organizing Map (SOM) method (standard settings) implemented in the Multiple Experiment Viewer (MeV) data analysis application. Numbers 1 to 6 refer to clusters 1 to 6. For the complete data, see **Supplemental Table S1**.

In summary, dipeptide levels respond to changes in environmental conditions. The exact response patterns could be linked to dipeptide amino acid composition.

### Level of acidic dipeptides correlates with autophagy-related transcripts across the stress time-course

To look for putative protein candidates responsible for the accumulation of acidic dipeptides, we correlated dipeptide levels (average of all dipeptides in cluster 4, see above and **Figure 1**) with the available transcript data from the same stress time-course [10]. We selected the 100 top correlated transcripts (referred to as the ‘positive list’) and looked at the functionality of the encoded proteins using the STRING database [15]. Remarkably, the positive list was enriched with transcripts encoding proteins associated with autophagy and proteolysis (Figure 2A-B; Supplementary Table S2). This included autophagy-associated proteins 8 (ATG8C, ATG8H, and ATG8F), 12 (ATG12A), 13 (ATG13) and 18 (ATG18G), multiple ubiquitin-protein ligases, cathepsin B-like cysteine protease (AtCathB2), serine carboxypeptidase S28, cysteine-type peptidase RD21, and aspartyl aminopeptidase (AT5G60160). Interestingly, three of the four proteases from the positive list are located in the vacuole [13], which is also the final destination of proteins captured by autophagosomes. Moreover, AtCathB2 has an experimentally tested dipeptidyl peptidase activity [16].

**Figure 2.**
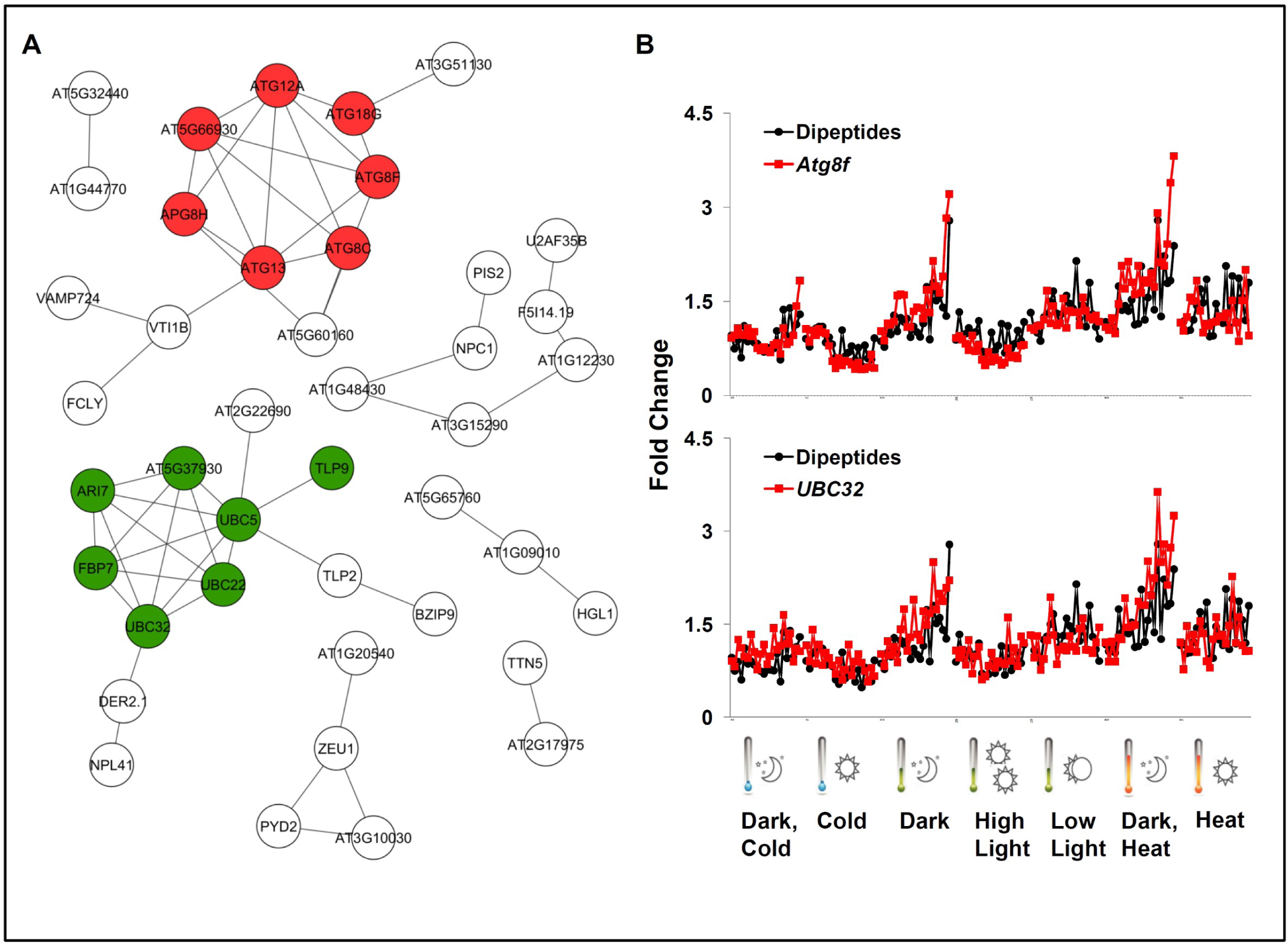
Transcripts co-related with acidic dipeptides in the stress time-course experiment. **(A)** Proteins encoded by the 100 top positively co-related transcripts were queried against the STRING protein-protein interaction database [15] and visualized using Cytoscape [34]. The edges represent medium confidence interactions (SCORE>0.4) based on experimental, database, and literature evidence. Red shading indicates proteins associated with autophagy and green shading with protein ubiquitination. Unconnected nodes are not displayed. **(B)** Accumulation of the acidic dipeptides (average of all dipeptides in cluster 4) and two representative transcripts from the positive list of the time-course stress experiment. Data are expressed as fold changes calculated in comparison to the control treatment (paired control). For the complete data, see **Supplemental Table S2**.

In summary, we showed that acidic dipeptides accumulate under conditions of darkness and heat stress **(see Figure 1)**, both of which are known to induce autophagy and proteolysis, and on par dipeptide levels are tightly correlated with the levels of transcripts encoding associated autophagy proteins, ubiquitin-protein ligases, and vacuolar proteases including dipeptidyl peptidases.

### Dipeptide accumulation during recovery from heat stress depends on autophagy

To examine whether the accumulation of dipeptide characteristics for dark and heat stress conditions (see above) depends on autophagy, we next employed a genetic approach. To this end, two different single (*atg18* and *nbr1-2*) and two double (*atg4(4a/4b*), *nbr1-2/atg5*) mutant lines defective in core and selective autophagy were grown on media plates, and 5-day-old seedlings were subjected to a two-step heat treatment established by [17] (see Materials and Methods). Samples were harvested two days after the heat treatment (within the recovery phase). Our experimental design was based on recent findings, which demonstrated a peak of autophagy at 2-3 days following heat stress treatment [17]. Collected seedlings were used for untargeted LC-MS-based metabolomics analysis, and a total of 365 metabolic features comprising dipeptides, primary and secondary metabolites, and lipids were putatively annotated to a compound using an in-house reference compound library [18].

Statistical analysis of the 57 measured dipeptides revealed striking differences between wild-type and mutant lines of Arabidopsis. While under control conditions, dipeptides were overall more abundant in all the four mutant lines when compared to the wild type; the opposite was observed in the heat stress recovery samples. That is because heat induced a significant increase in the dipeptide levels in the wild type, but no such increase was measured in the autophagy mutants. In contrast to the stress time-course experiment, 52 of the measured dipeptides responded in a similar fashion **(Figure 3 and Supplemental Figure S1)**. There were, however, five exceptions: Asp-Thr, Glu-Glu, Glu-Gln, Ala-Thr, and Val-Lys **(Supplemental Figure S1 and Figure S2)**. Whilst levels of both Asp-Thr and Val-Lys were significantly (Student’s *t*-test *p*-value <0.05) decreased by stress in the wild-type plants, Glu-Glu was not affected, and the levels of Glu-Gln and Ala-Thr significantly increased in both wild-type and mutant plants (**Supplemental Figure S2)**.

**Figure 3.**
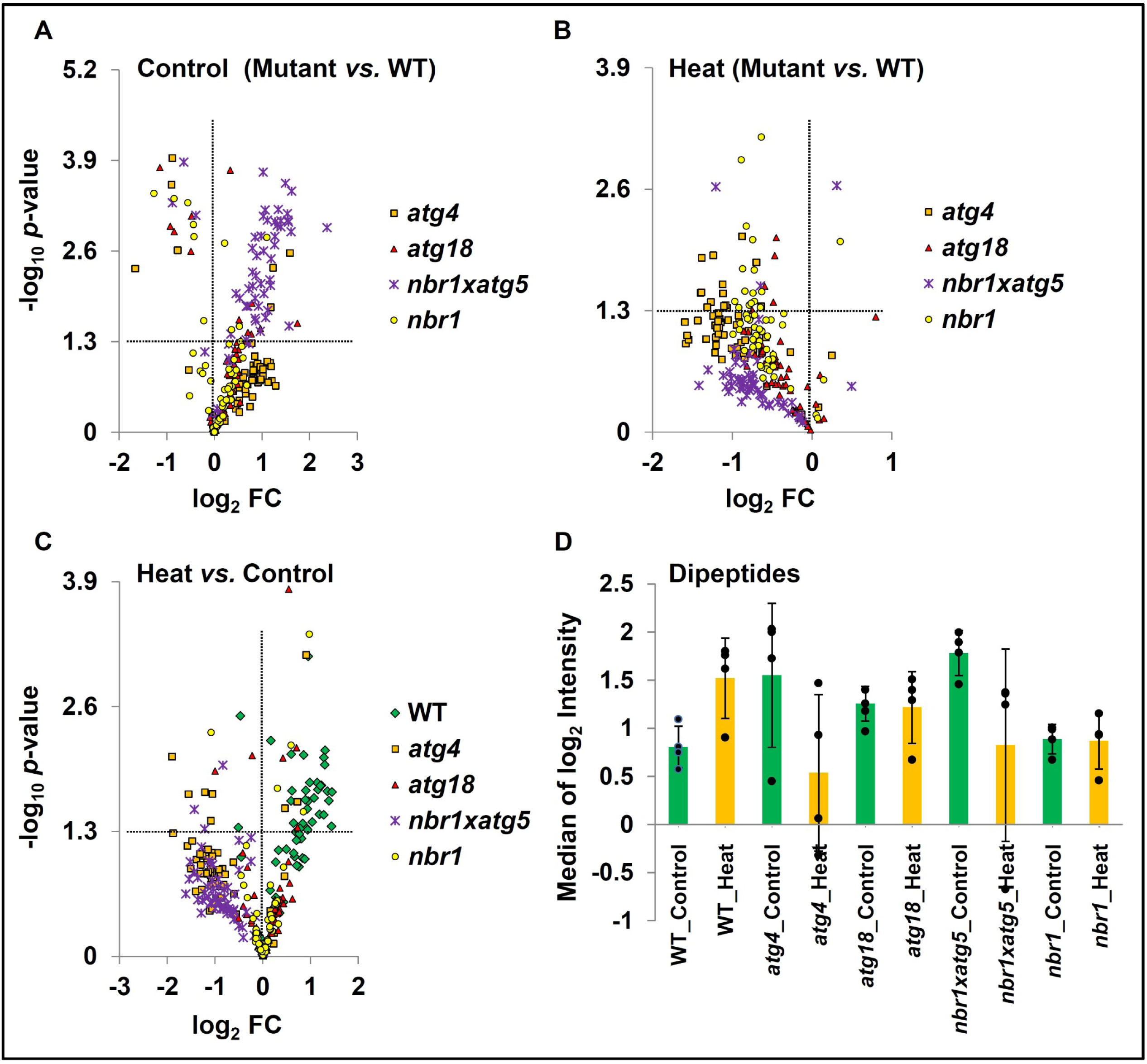
Dipeptide accumulation measured during heat stress recovery depends on the autophagy. Volcano plot representation of the log_2_ fold changes (axis x) in dipeptide levels in the function of –log_10_ of p-value (axis y). *P*-value was calculated using unpaired Student’s *t*-test with two-tailed distribution assuming unequal variance; *n*=4 of biological replicates. **(A)** Between wild-type (WT) and autophagy-deficient mutants grown under control conditions. **(B)** Between wild-type (WT) and autophagy-deficient mutants measured two days into recovery from heat treatment. **(C)** Between heat and control samples for wild-type (WT) and autophagy-deficient mutants. **(D)** The median of all dipeptides was used to calculate the average and standard deviation (SD) across mutants and treatments. Note that data for individual dipeptides used to calculate the median were expressed as log_2_ of normalized LC-MS intensities. **(A-C)** Horizontal dash line indicates *p*-value of 0.05. For the complete data, see **Supplemental Table S3**.

In conclusion, dipeptide accumulation measured during heat stress recovery depends on autophagy.

### Metabolite profiling of autophagy-deficient mutants reveals significant metabolome alterations

The canonical role of autophagy is the recycling of cellular constituents, including nutrients. Thus, not surprisingly, autophagy-deficient mutants are characterized by significant alterations in the metabolome under both control and stress conditions [12,19,20]. Changes in the accumulation of amino acids, sugars, lipids, and secondary metabolites have been reported and were shown to be strongly dependent on the plant’s developmental stage and growth conditions. For instance, etiolated seedlings of the autophagy-deficient mutant *atg5* are characterized by reduced levels of amino acids and glucosinolates, while the levels of triacylglycerols, fatty acids, and sugars increased [12]. By contrast, mature rosette leaves of the *atg5* mutant were shown to accumulate more amino acids but fewer sugars and anthocyanins under both control and nitrate-limiting conditions [19].

Herein, we present primary and secondary metabolites and lipid data for Arabidopsis seedlings grown under control conditions and subjected to heat stress, followed by a recovery (**Supplemental Table S3)**. Notably, all of the four tested autophagy mutants were characterized by significant alterations in their metabolome, already under control conditions **(Figure 4, Supplemental Figure S3)**. Amino acids and flavonols were overall lower in the mutants, irrespective of the treatment, whereas phospholipids, lysophospholipids, nucleotides, and cofactors NADH, NADP^+^, and FAD^+^ were higher, but only under control conditions. While heat stress induced nucleotide accumulation in the wild-type plants, no such accumulation was measured in the autophagy mutants, consistent with the involvement of autophagy in nucleic acid degradation. Another interesting observation relates to the difference in the accumulation of triacylglycerols (TAGs) in *nbr1* plants in comparison to the wild type and the other autophagy mutants. While TAGs 42-50 (the number relates to the total carbon length of fatty acid chains) were overall lower in all mutants in comparison to the wild type, the much more abundant TAGs 50-60 were significantly decreased in the *nbr1* mutant only. In contrast to the *atg4, atg18*, and *atg5* mutants, *nbr1* is specifically defective in selective autophagy **(Supplemental Figure S4)**. The differential accumulation of TAGs in *nbr1* plants is therefore an exciting finding for a follow-up analysis.

**Figure 4.**
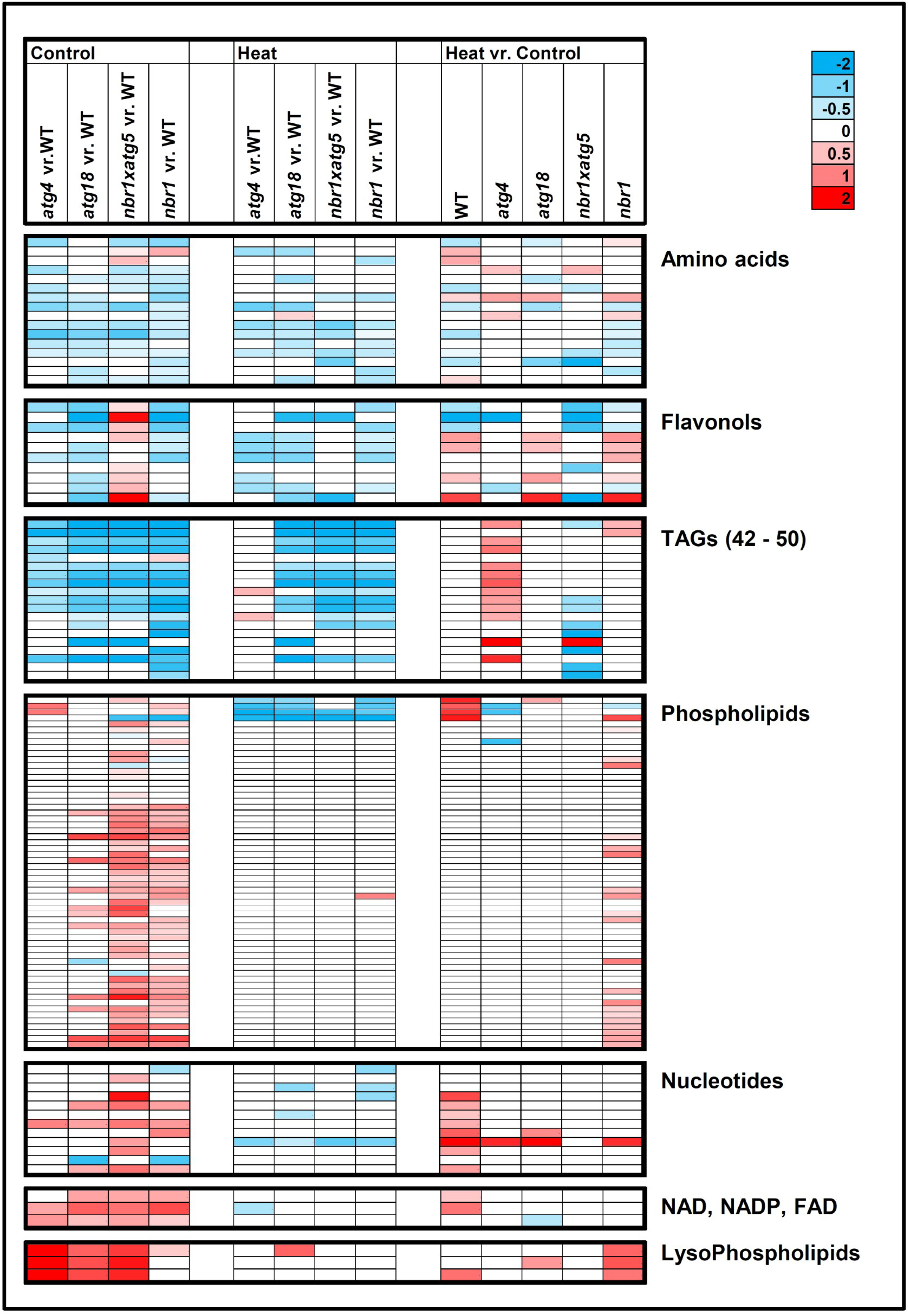
Metabolomic alterations in autophagy-deficient mutants. The heat map represents the significant (*P*-value < 0.05) log_2_ fold changes of the selected metabolites (for the full dataset, see **Supplemental Table S3**). *P*-value was calculated using unpaired Student’s *t*-test with two-tailed distribution assuming unequal variance; *<u>n</u>*=4 biological replicates.

In summary, the results presented demonstrate the essential role of autophagy for the regulation of plant metabolism.

## Discussion

### Dipeptide accumulation depends on autophagy

The prime role of autophagy is the recycling of macromolecules. Proteins captured by autophagosomes are delivered to the vacuoles, where they are cleaved to amino acids, which in turn get transported back to the cytosol by amino acid transporters situated in a vacuolar membrane. Herein, we demonstrate that autophagy, in addition to amino acid recycling, is also responsible for the increased accumulation of dipeptides in recovery from heat stress **(Figures 2-3)**. In support of our results, vacuolar peptidases with dipeptidyl peptidase activity, e.g., AtCathB2; At1g02305 and serine carboxypeptidase S28, At5g60360 [13,16], as well as vacuolar dipeptide transporters, e.g., PTR2; AT2G02040 [13,21], have been reported.

Dipeptides can be further cleaved to single amino acids, and as such, represent the intermediates of proteolysis. However, proteogenic dipeptides have also been reported to function as small-molecule regulators. In addition to the already given examples (see Introduction), dipeptides were shown, for instance, to activate the p38MAPK-Smad3 signaling pathway in the rare chronic myelogenous leukemia (CML) stem cell population, and in line pharmacological inhibition of dipeptide uptake inhibited CML stem cell activity [22]. Dipeptides were also reproducibly measured in the Arabidopsis root exudates, downstream of MPK3/6 signaling, suggesting a role in plant-microbe and plant-plant communication [23]. An inhibitory effect of five different dipeptides on maize root growth was reported previously, supporting such a role [24], while phenylalanine-containing dipeptides accumulate in wheat upon treatment with deoxynivalenol, the major toxin and virulence factor of the fungal pathogen *Fusarium graminearum* [25].

Interestingly, in the context of our findings, a single dipeptide Tyr-Ala, but neither tyrosine nor alanine, was shown to significantly enhance the lifespan and healthspan of the model nematode *Caenorhabditis elegans* [26]. Tyr-Ala activity was associated with the upregulation of stress-related proteins, ROS (reactive oxygen species) scavenging enzymes, and chaperones. In agreement, Tyr-Ala effects were particularly pronounced under heat and oxidative stress conditions, reminiscent of the dipeptide accumulation reported in our study. Moreover, we previously demonstrated that Tyr-Asp binds to a glycolytic enzyme GAPDHs [6], and inhibition of GAPDH activity shifts glycolytic flux towards the pentose phosphate pathway and production of NADPH, the primary source of reducing power for biosynthesis and detoxification of ROS during stress [27]. Moreover, dipeptides were shown to promote protein folding, which is important under stress conditions [28]. Based on the presented results and on literature knowledge, we speculate that the dipeptide accumulation observed under heat conditions, and related to autophagy, serves to alleviate oxidative stress and protein damage.

### Note on the specificity of dipeptide accumulation under different environmental regimes

An important question about dipeptide biogenesis is specificity. Analysis of the stress time-course experiment revealed that dipeptide levels are stress-responsive, but what’s more intriguing is that not all the dipeptides respond in the same manner **(Figure 1)**. Amino acid composition, such as the presence of an acidic or branch-chain amino acid, defined groups of dipeptides sharing similar stress behavior, but the different groups behaved differently. On par with our results, a recent metabolomics analysis of different root cell types revealed preferential accumulation of Leu and Ile containing dipeptides in the endodermis and epidermis [29]. By contrast, heat treatment of the Arabidopsis seedlings **(Figure 3**, referred to as seedling experiment) resulted in the bulk accumulation of dipeptides, with no apparent specificity. Of the 57 dipeptides, 52 were characterized by similar accumulation patterns. The five remaining dipeptides could not be easily clustered based on either their amino acid composition or response pattern.

However, both the time-course and seedling experiments exploited heat stress; the regimes were very different. While the seedling experiment is representative of a severe stress regime resulting from a combination of *in vitro* growth conditions, young plants, and severe heat treatment (37 °C and 44 °C), the time-course experiment was performed using soil-grown, mature plants subjected to prolonged but mild heat conditions (32 °C), which do not threaten plant survival. Moreover, in the seedling experiment, rather than looking at heat stress *per se*, samples were harvested in the recovery-phase after the heat stress. However, it was demonstrated that heat activates autophagy and increases the number of autophagosomes and autophagic bodies during the recovery phase [17]. Recycling of the stress-associated proteins, such as heat shock proteins, is considered an essential component of the stress-resetting mechanism. Bulk accumulation patterns that are characteristic for the seedling experiment could thus be explained by the rapid turnover of multiple stress proteins without the preference for particular dipeptide motifs. By contrast, prolonged and mild heat stress would favor more selective recycling, leading to the specificity in dipeptide accumulation. This is a hypothesis that needs to be addressed in future studies.

## Materials and Methods

### Dipeptide and transcript data: time-course stress experiment

Time-course experiment [10]: Briefly, mature Arabidopsis plants grown in soil at 21 °C and in standard light (150 µE m^−2^ s^−1^) were either kept at this condition or transferred into seven different environments varying in temperature and/or light intensity. More specifically, the seven conditions were as follows; low light (75 µE m^−2^ s^−1^), darkness, high light (300 µE m^−2^ s^−1^), heat (32 °C, 150 µE m^−2^ s^−1^), cold (4° C, 85 µE m^−2^ s^−1^), cold and darkness, and heat and darkness. Samples were harvested at 22 different time points ranging from 5 minutes to approximately 21 hours after the transfer. Raw metabolite LC-MS data from the time-course stress experiment were retrieved from [11]. We used the retrieved dataset to annotate dipeptides by matching metabolic features, previously assigned as unknown, with a dipeptide reference compound library. Data were subjected to day-normalization and sample-median-normalization. Dipeptide annotation was made, allowing 0.15 RT and 5 ppm deviation from the dipeptide reference compound library comprising 331 dipeptides. Transcript data used for co-expression analysis were retrieved from [10].

### Plant materials and growth conditions

Seeds of *A. thaliana* ecotype Col-0 and the mutant or transgenic lines *nbr1-2* [30,31], *nbr1-2/atg5* [30,31], (*atg4(4a/4b)* [32], and *atg18a-2* [33] were surface-sterilized and sown in Petri dishes containing Murashige-Skoog (MS) agar medium supplemented with 1% (w/v) sucrose. They were stratified at 4 °C in darkness for 2 days, and seedlings were grown in growth chambers providing 16 h light/8 h dark cycles at 22 °C.

### Heat stress experiment

Heat stress (also referred to as priming) was applied in two steps [17]. Five-day-old seedlings were subjected to an incubation (with an incubator) at 37 °C for 1.5 h, then at 22 °C for a 1.5□h recovery period, followed by 45□min of heat stress at 44□ °C (in a hot water bath). After the treatment, the seedlings were transferred to normal growth conditions (16 h light/8 h dark cycles) at 22 °C for 2 days (recovery phase), during which samples (whole seedlings) were harvested for analyses.

### Metabolite extraction

The extraction protocol was adapted and modified from [18]. Proteins and metabolites were extracted using a methyl-*tert*-butyl ether (MTBE)/methanol/water solvent system, which separates molecules into pellets (proteins), organics (lipids), and an aqueous phase (primary and secondary metabolites). Equal volumes of the polar and lipid fractions were dried in a centrifugal evaporator and stored at –80 °C until further processing.

### LC-MS metabolomics

The dried aqueous phase was measured using ultra-performance liquid chromatography coupled to an Exactive mass spectrometer (Thermo Fisher Scientific) in positive and negative ionization modes, as described in [18]. A 2-μl sample (the dried aqueous fraction was resuspended in 200 μl of UHPLC-grade water) was loaded per injection. Similarly, the dried organic phase was measured using ultra-performance liquid chromatography coupled to an Exactive mass spectrometer (Thermo Fisher Scientific) in positive and negative ionization modes, as described in [18]. A 2-μl sample (the dried organic fraction was resuspended in 200 μl of UPLC-grade 7:3 acetonitrile:isopropanol) was loaded per injection.

### Data pre-processing: LC-MS metabolite data

The LC-MS data were processed using Expressionist Refiner MS 11.0 (Genedata AG, Basel, Switzerland). Settings were as follows: chromatogram alignment (RT search interval, 0.5 min), peak detection (summation window, 0.09 min; minimum peak size, 0.03 min; gap/peak ratio, 50%; smoothing window, 5 points; center computation by intensity-weighted method with threshold at 70%; boundary determination using inflection points), isotope clustering (RT tolerance at 0.015 min, m/z tolerance 5 ppm, allowed charges 1–5), filtering for a single peak not assigned to an isotope cluster, adduct detection, and cluster grouping (RT tolerance 0.05 min, m/z tolerance 5 ppm, maximum intensity of side adduct 100000%). All metabolite clusters were matched to in-house libraries of reference compounds, allowing a 10-ppm mass and dynamic retention time deviation (maximum 0.1 min). All annotations are putative.

### Data analysis

Raw metabolites’ intensities were normalized to the median of chromatogram intensity in order to correct for variation in sample extraction and measurement. Prior statistical analysis data were subjected to log_2_ transformation. *P*-value was calculated using unpaired Student’s *t*-test with two-tailed distribution; n=4 of biological replicates. Heat stress experiment was replicated with individual plates grown in one experiment. The data are presented in **Supplemental Tables S3**.

## Supporting information

Figyres S1 to S4

## Supplemental figure legends

**Supplemental Figure S1**. Heat map of log_2_ fold change of dipeptides measured in heat recovery in comparison to control conditions. The data for all samples are given. Clustering was performed using ClustVis tool (https://biit.cs.ut.ee/clustvis/) and default settings. Please see Supplemental Table S3.

**Supplemental Figure S2**. The median of the fold change of all dipeptides measured in the heat versus control conditions, separately in the wild type and the mutants, was used to calculate Pearson correlation with all the dipeptides individually. For five of the measured dipeptides (indicated by an orange shading), the Pearson correlation was below 0.5, indicating a difference in response pattern.

**Supplemental Figure S3**. PCA plot prepared using all annotated metabolites in wild type (WT) and mutants grown under control conditions using ClustVis tool (https://biit.cs.ut.ee/clustvis/) and default settings. Please see Supplemental Table S3.

**Supplemental Figure S4. TAG accumulation in the wild type and in the *nbr1* mutant measured under control conditions**. Data are expressed as mean of the median normalized intensity. *P*-value was calculated using unpaired Student’s *t*-test with two-tailed distribution assuming unequal variance; *n*=4 of biological replicates. Asterisks indicate significance (*P*-value < 0.05).

## Supplemental Tables

**Supplementary Table S1**. Dipeptide data for Figure 1. The fold change in comparison to the paired control is given. D, darkness; 4, cold; 30, heat; LL, low light; HL, high light. The time is given in minutes. For example, 4D5: cold and dark stress, five minutes after the stress onset.

**Supplementary Table S2**. Transcript data for Figure 2. The fold change in comparison to the paired control is given. D, darkness; 4, cold; 30, heat; LL, low light; HL, high light. The time is given in minutes. For example, 4D5: cold and dark stress, five minutes after the stress onset. Pearson correlation (column D) was calculated using the average response of dipeptides from cluster 4 (refer Supplementary Table S1).

**Supplementary Table S3**. Normalized and log_2_ transformed metabolite data for Figure 3 and Figure 4. Raw metabolite intensities were normalized to the median of chromatogram intensity in order to correct for variation in sample extraction and measurement. Prior statistical analysis data were subjected to log_2_ transformation.

## Acknowledgements

We thank Anne Michaelis for running the metabolomics samples and Prof. Alisdair R. Fernie for his valuable scientific input. Salma Balazadeh thanks the Deutsche Forschungsgemeinschaft for the funding (CRC 973).

## Author Contributions

A.S. and V.P.K. conceived the experiments and wrote the manuscript. V.P.K. performed experiments within the DFG-funded research project of S.B. M.W. performed metabolic extractions. A.S. analyzed the data.

## Conflicts of Interest

The authors declare no conflict of interest.

## References

1. Du, F.Y.; Navarro-Garcia, F.; Xia, Z.X.; Tasaki, T.; Varshavsky, A. Pairs of dipeptides synergistically activate the binding of substrate by ubiquitin ligase through dissociation of its autoinhibitory domain. Proceedings of the National Academy of Sciences of the United States of America 2002, 99, 14110–14115, doi:DOI 10.1073/pnas.172527399.

2. Takagi, H.; Shiomi, H.; Ueda, H.; Amano, H. Morphine-Like Analgesia by a New Dipeptide, L-Tyrosyl-L-Arginine (Kyotorphin) and Its Analog. European Journal of Pharmacology 1979, 55, 109–111, doi:Doi 10.1016/0014-2999(79)90154-7.

3. Ueda, H.; Yoshihara, Y.; Fukushima, N.; Shiomi, H.; Nakamura, A.; Takagi, H. Kyotorphin (Tyrosine-Arginine) Synthetase in Rat-Brain Synaptosomes. Journal of Biological Chemistry 1987, 262, 8165–8173.

4. Mizushige, T.; Kanegawa, N.; Yamada, A.; Ota, A.; Kanamoto, R.; Ohinata, K. Aromatic amino acid-leucine dipeptides exhibit anxiolytic-like activity in young mice. Neuroscience Letters 2013, 543, 126–129, doi:10.1016/j.neulet.2013.03.043.

5. Veyel, D.; Kierszniowska, S.; Kosmacz, M.; Sokolowska, E.M.; Michaelis, A.; Luzarowski, M.; Szlachetko, J.; Willmitzer, L.; Skirycz, A. System-wide detection of protein-small molecule complexes suggests extensive metabolite regulation in plants. Scientific Reports 2017, 7, 42387, doi:10.1038/srep42387.

6. Veyel, D.; Sokolowska, E.M.; Moreno, J.C.; Kierszniowska, S.; Cichon, J.; Wojciechowska, I.; Luzarowski, M.; Kosmacz, M.; Szlachetko, J.; Gorka, M., et al. PROMIS, global analysis of PROtein-Metabolite Interactions using Size separation in Arabidopsis thaliana. The Journal of Biological Chemistry 2018, 10.1074/jbc.RA118.003351, doi:10.1074/jbc.RA118.003351.

7. Zaffagnini, M.; Fermani, S.; Costa, A.; Lemaire, S.D.; Trost, P. Plant cytoplasmic GAPDH: redox post-translational modifications and moonlighting properties. Frontiers in Plant Science 2013, 4, 450, doi:10.3389/fpls.2013.00450.

8. Kenny, A.J.; Booth, A.G.; George, S.G.; Ingram, J.; Kershaw, D.; Wood, E.J.; Young, A.R. Dipeptidyl peptidase IV, a kidney brush-border serine peptidase. The Biochemical Journal 1976, 157, 169–182, doi:10.1042/bj1570169.

9. Kumar, S.; Kaur, A.; Chattopadhyay, B.; Bachhawat, A.K. Defining the cytosolic pathway of glutathione degradation in Arabidopsis thaliana: role of the ChaC/GCG family of gamma-glutamyl cyclotransferases as glutathione-degrading enzymes and AtLAP1 as the Cys-Gly peptidase. The Biochemical Journal 2015, 468, 73–85, doi:10.1042/BJ20141154.

10. Caldana, C.; Degenkolbe, T.; Cuadros-Inostroza, A.; Klie, S.; Sulpice, R.; Leisse, A.; Steinhauser, D.; Fernie, A.R.; Willmitzer, L.; Hannah, M.A. High-density kinetic analysis of the metabolomic and transcriptomic response of Arabidopsis to eight environmental conditions. Plant J 2011, 67, 869–884, doi:10.1111/j.1365-313X.2011.04640.x.

11. Wu, S.; Tohge, T.; Cuadros-Inostroza, A.; Tong, H.; Tenenboim, H.; Kooke, R.; Meret, M.; Keurentjes, J.B.; Nikoloski, Z.; Fernie, A.R., et al. Mapping the Arabidopsis Metabolic Landscape by Untargeted Metabolomics at Different Environmental Conditions. Mol Plant 2018, 11, 118–134, doi:10.1016/j.molp.2017.08.012.

12. Avin-Wittenberg, T.; Bajdzienko, K.; Wittenberg, G.; Alseekh, S.; Tohge, T.; Bock, R.; Giavalisco, P.; Fernie, A.R. Global analysis of the role of autophagy in cellular metabolism and energy homeostasis in Arabidopsis seedlings under carbon starvation. The Plant Cell 2015, 27, 306–322, doi:10.1105/tpc.114.134205.

13. Carter, C.; Pan, S.; Zouhar, J.; Avila, E.L.; Girke, T.; Raikhel, N.V. The vegetative vacuole proteome of Arabidopsis thaliana reveals predicted and unexpected proteins. The Plant Cell 2004, 16, 3285–3303, doi:10.1105/tpc.104.027078.

14. Marion, J.; Le Bars, R.; Besse, L.; Batoko, H.; Satiat-Jeunemaitre, B. Multiscale and Multimodal Approaches to Study Autophagy in Model Plants. Cells 2018, 7, doi:10.3390/cells7010005.

15. von Mering, C.; Huynen, M.; Jaeggi, D.; Schmidt, S.; Bork, P.; Snel, B. STRING: a database of predicted functional associations between proteins. Nucleic Acids Res 2003, 31, 258–261.

16. Porodko, A.; Cirnski, A.; Petrov, D.; Raab, T.; Paireder, M.; Mayer, B.; Maresch, D.; Nika, L.; Biniossek, M.L.; Gallois, P., et al. The two cathepsin B-like proteases of Arabidopsis thaliana are closely related enzymes with discrete endopeptidase and carboxydipeptidase activities. Biological Chemistry 2018, 399, 1223–1235, doi:10.1515/hsz-2018-0186.

17. Sedaghatmehr, M.; Thirumalaikumar, V.P.; Kamranfar, I.; Marmagne, A.; Masclaux-Daubresse, C.; Balazadeh, S. A regulatory role of autophagy for resetting the memory of heat stress in plants. Plant, Cell & Environment 2019, 42, 1054–1064, doi:10.1111/pce.13426.

18. Giavalisco, P.; Li, Y.; Matthes, A.; Eckhardt, A.; Hubberten, H.M.; Hesse, H.; Segu, S.; Hummel, J.; Kohl, K.; Willmitzer, L. Elemental formula annotation of polar and lipophilic metabolites using (13) C, (15) N and (34) S isotope labelling, in combination with high-resolution mass spectrometry. The Plant 1. Journal : for cell and molecular biology 2011, 68, 364–376, doi:10.1111/j.1365-313X.2011.04682.x.

19. Masclaux-Daubresse, C.; Clement, G.; Anne, P.; Routaboul, J.M.; Guiboileau, A.; Soulay, F.; Shirasu, K.; Yoshimoto, K. Stitching together the Multiple Dimensions of Autophagy Using Metabolomics and Transcriptomics Reveals Impacts on Metabolism, Development, and Plant Responses to the Environment in Arabidopsis. The Plant Cell 2014, 26, 1857–1877, doi:10.1105/tpc.114.124677.

20. McLoughlin, F.; Augustine, R.C.; Marshall, R.S.; Li, F.; Kirkpatrick, L.D.; Otegui, M.S.; Vierstra, R.D. Maize multi-omics reveal roles for autophagic recycling in proteome remodelling and lipid turnover. Nature Plants 2018, 4, 1056–1070, doi:10.1038/s41477-018-0299-2.

21. Chiang, C.S.; Stacey, G.; Tsay, Y.F. Mechanisms and functional properties of two peptide transporters, AtPTR2 and fPTR2. The Journal of Biological Chemistry 2004, 279, 30150–30157, doi:10.1074/jbc.M405192200.

22. Naka, K.; Jomen, Y.; Ishihara, K.; Kim, J.; Ishimoto, T.; Bae, E.J.; Mohney, R.P.; Stirdivant, S.M.; Oshima, H.; Oshima, M., et al. Dipeptide species regulate p38MAPK-Smad3 signalling to maintain chronic myelogenous leukaemia stem cells. Nature Communications 2015, 6, 8039, doi:10.1038/ncomms9039.

23. Strehmel, N.; Hoehenwarter, W.; Monchgesang, S.; Majovsky, P.; Kruger, S.; Scheel, D.; Lee, J. Stress-Related Mitogen-Activated Protein Kinases Stimulate the Accumulation of Small Molecules and Proteins in Arabidopsis thaliana Root Exudates. Front Plant Sci 2017, 8, doi:Artn 1292 10.3389/Fpls.2017.01292.

24. Liu, D.L.Y.; Christians, N.E. Isolation and Identification of Root-Inhibiting Compounds from Corn Gluten Hydrolysate. J Plant Growth Regul 1994, 13, 227–230, doi:Doi 10.1007/Bf00226041.

25. Doppler, M.; Kluger, B.; Bueschl, C.; Steiner, B.; Buerstmayr, H.; Lemmens, M.; Krska, R.; Adam, G.; Schuhmacher, R. Stable Isotope-Assisted Plant Metabolomics: Investigation of Phenylalanine-Related Metabolic Response in Wheat Upon Treatment With the Fusarium Virulence Factor Deoxynivalenol. Frontiers in Plant Science 2019, 10, doi:Artn 1137 10.3389/Fpls.2019.01137.

26. Zhang, Z.; Zhao, Y.; Wang, X.; Lin, R.; Zhang, Y.; Ma, H.; Guo, Y.; Xu, L.; Zhao, B. The novel dipeptide Tyr-Ala (TA) significantly enhances the lifespan and healthspan of Caenorhabditis elegans. Food Funct 2016, 7, 1975–1984, doi:10.1039/c5fo01302j.

27. Dick, T.P.; Ralser, M. Metabolic Remodeling in Times of Stress: Who Shoots Faster than His Shadow? Molecular Cell 2015, 59, 519–521, doi:10.1016/j.molcel.2015.08.002.

28. Saini, S.K.; Ostermeir, K.; Ramnarayan, V.R.; Schuster, H.; Zacharias, M.; Springer, S. Dipeptides promote folding and peptide binding of MHC class I molecules. Proceedings of the National Academy of Sciences of the United States of America 2013, 110, 15383–15388, doi:10.1073/pnas.1308672110.

29. Moussaieff, A.; Rogachev, I.; Brodsky, L.; Malitsky, S.; Toal, T.W.; Belcher, H.; Yativ, M.; Brady, S.M.; Benfey, P.N.; Aharoni, A. High-resolution metabolic mapping of cell types in plant roots. Proceedings of the National Academy of Sciences of the United States of America 2013, 110, E1232–E1241, doi:10.1073/pnas.1302019110.

30. Hafren, A.; Hofius, D. NBR1-mediated antiviral xenophagy in plant immunity. Autophagy 2017, 13, 2000–2001, doi:10.1080/15548627.2017.1339005.

31. Hafren, A.; Macia, J.L.; Love, A.J.; Milner, J.J.; Drucker, M.; Hofius, D. Selective autophagy limits cauliflower mosaic virus infection by NBR1-mediated targeting of viral capsid protein and particles. Proceedings of the National Academy of Sciences of the United States of America 2017, 114, E2026–E2035, doi:10.1073/pnas.1610687114.

32. Chung, T.; Phillips, A.R.; Vierstra, R.D. ATG8 lipidation and ATG8-mediated autophagy in Arabidopsis require ATG12 expressed from the differentially controlled ATG12A AND ATG12B loci. The Plant Journal : for cell and molecular biology 2010, 62, 483–493, doi:10.1111/j.1365-313X.2010.04166.x.

33. Shibata, M.; Oikawa, K.; Yoshimoto, K.; Kondo, M.; Mano, S.; Yamada, K.; Hayashi, M.; Sakamoto, W.; Ohsumi, Y.; Nishimura, M. Highly oxidized peroxisomes are selectively degraded via autophagy in Arabidopsis. The Plant Cell 2013, 25, 4967–4983, doi:10.1105/tpc.113.116947.

34. Shannon, P.; Markiel, A.; Ozier, O.; Baliga, N.S.; Wang, J.T.; Ramage, D.; Amin, N.; Schwikowski, B.; Ideker, T. Cytoscape: a software environment for integrated models of biomolecular interaction networks. Genome Research 2003, 13, 2498–2504, doi:10.1101/gr.1239303.

