## Supplementary material for "Autophagy is responsible for the accumulation of proteogenic dipeptides in response to heat stress in *Arabidopsis thaliana*": Figyres S1 to S4

### Supplemental figures

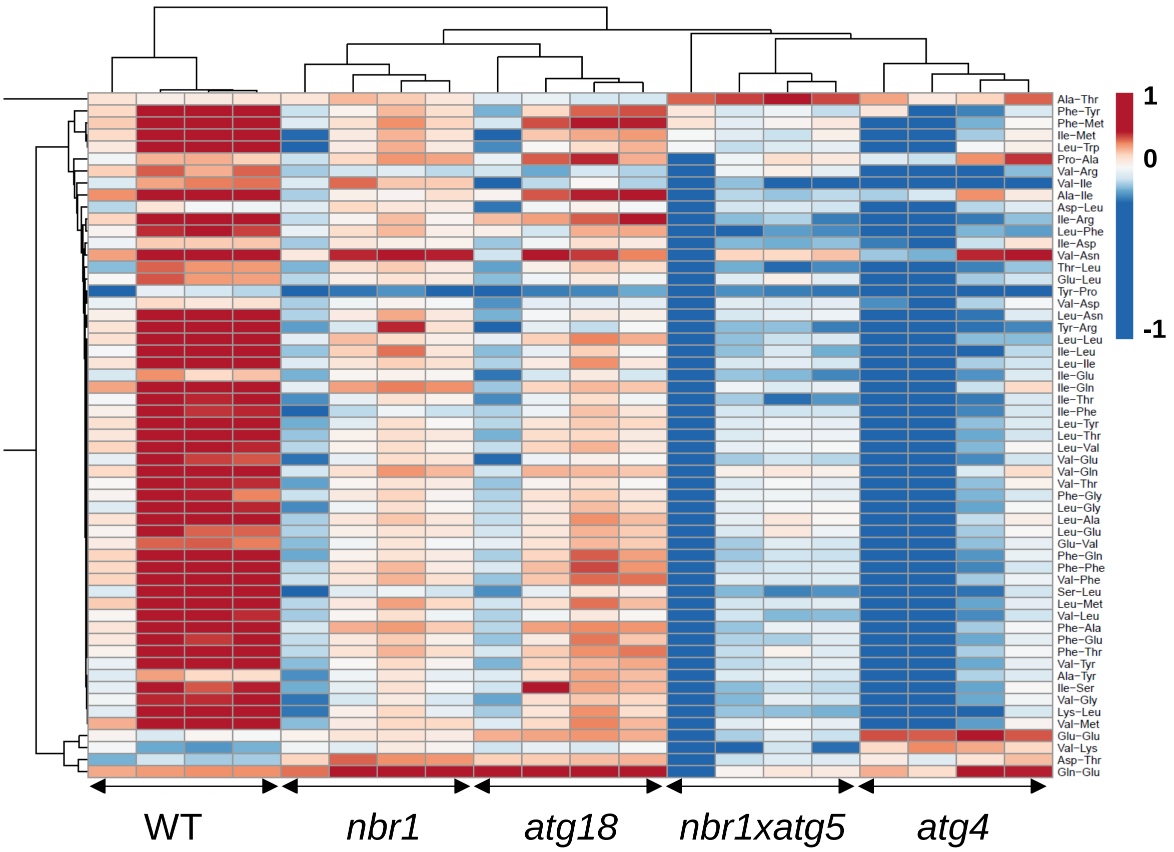


**Supplemental Figure S1.** Heat map of $log_2$ fold change of dipeptides measured in heat recovery in comparison to control conditions. Given are data for all samples. Clustering was performed using ClustVis tool (https://biit.cs.ut.ee/clustvis/) and default settings. Please see Supplemental Table S3.

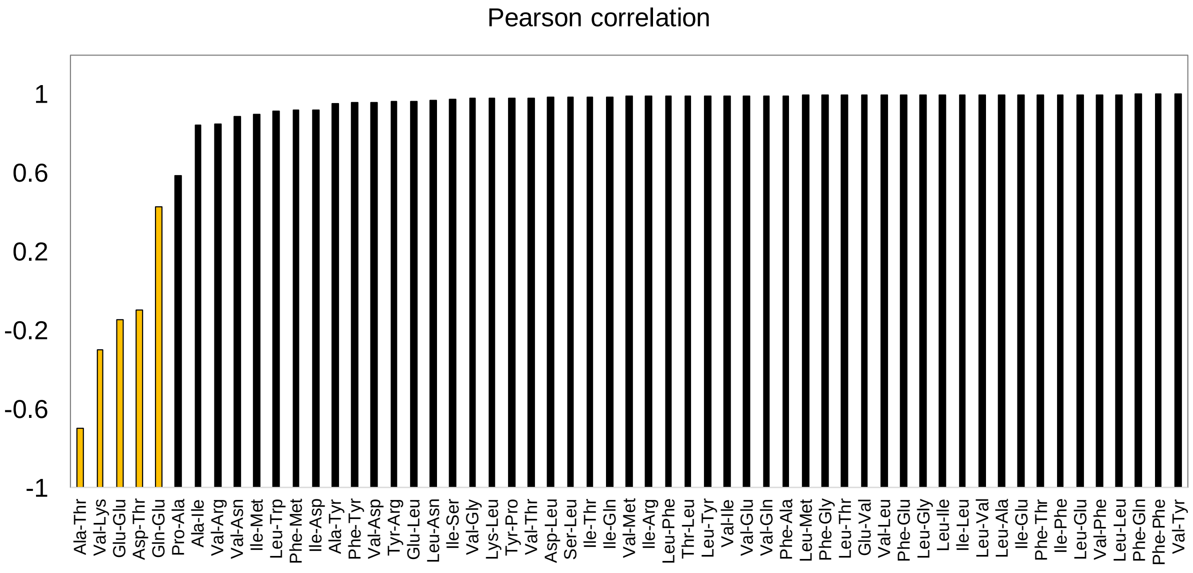


**Supplemental Figure S2.** Median of the fold change of all dipeptides measured in the heat versus control conditions, separately in the wild type and the mutants, was used to calculate Pearson correlation with all the dipeptides individually. For five of the measured dipeptides (indicated by an orange shading), the Pearson correlation was below 0.5, indicating a difference in response pattern.

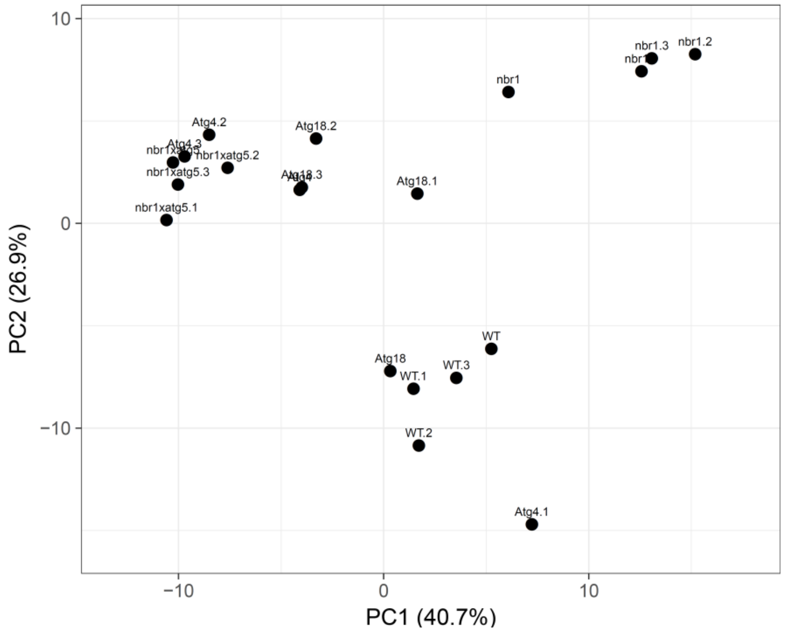


**Supplemental Figure S3.** PCA plot prepared using all annotated metabolites in wild type (WT) and mutants grown control conditions using ClustVis tool (https://biit.cs.ut.ee/clustvis/) and default settings. Please see Supplemental Table S3.

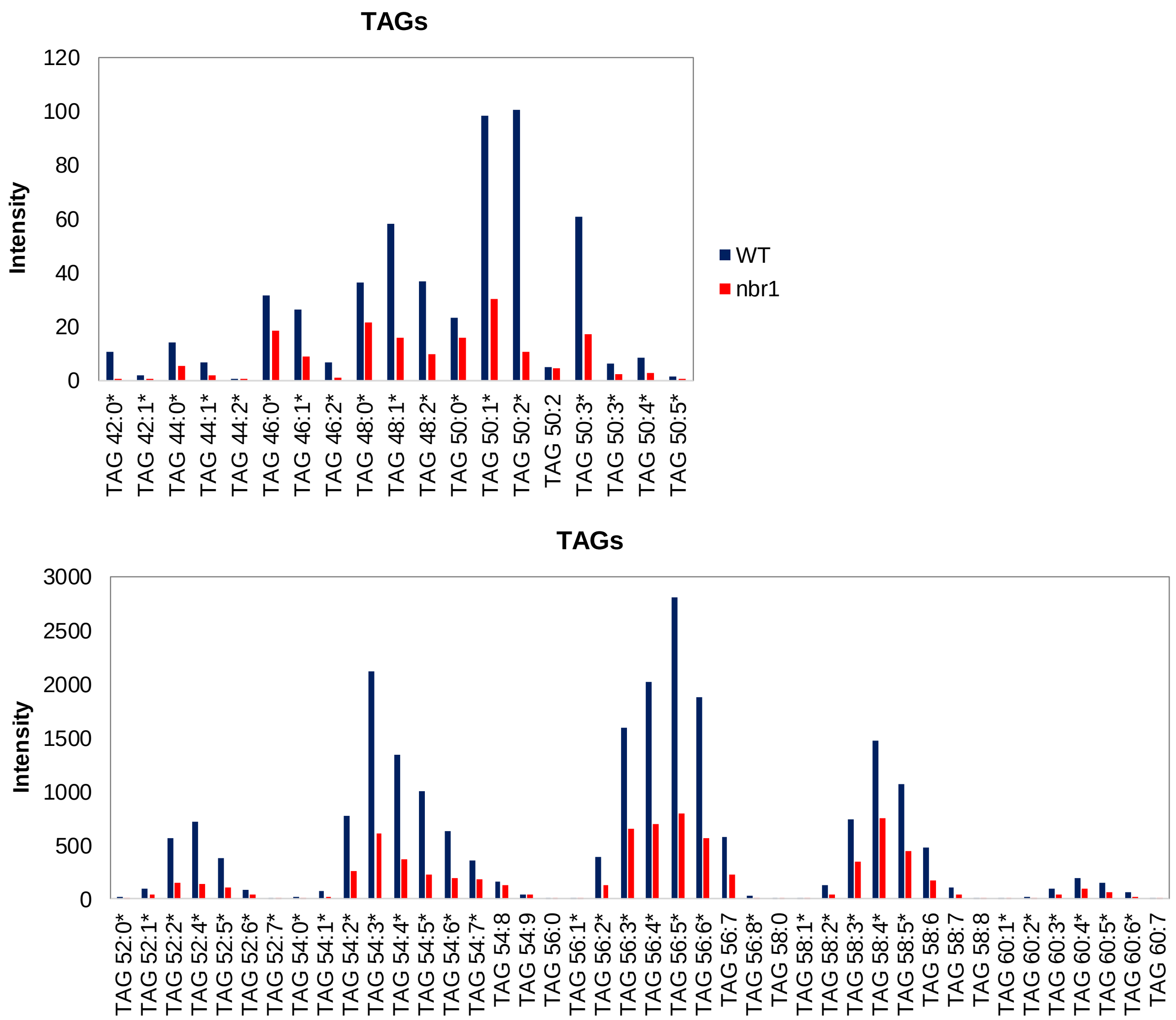


**Supplemental Figure S4. TAG accumulation in the wild type and in the *nbr1* mutant measured under control conditions.** Data are expressed as mean of the median normalized intensity. *P*-value was calculated using unpaired Student's *t*-test with two-tailed distribution assuming unequal variance; *n*=4 of biological replicates. Asteriks indicates significance (*P*-value < 0.05).
